# Intraspecific variation of two duckweed species influences response to microcystin-LR exposure

**DOI:** 10.1101/2023.06.04.543632

**Authors:** Lacey D. Rzodkiewicz, Martin M. Turcotte

## Abstract

Cyanotoxins produced by harmful cyanobacteria blooms can damage freshwater ecosystems and threaten human health. Floating macrophytes may be used as a means of biocontrol by limiting light and resources available to cyanobacteria. However, genetic variation in macrophyte sensitivity to cyanotoxins could influence their suitability as biocontrol agents. We investigated the influence of such intraspecific variation on the response of two rapidly growing duckweed species, *Lemna minor* and *Spirodela polyrhiza*, often used in nutrient and metal bioremediation. We assessed two biomarkers related to productivity (biomass and chlorophyll A production) and two related to fitness measures (population size and growth rate). Fifteen genetic lineages of each species were grown in media containing common cyanotoxin microcystin-LR at ecologically relevant concentrations or control media for a period of twelve days. Genotype identity had a strong impact on all biomarker responses. Microcystin concentration did impact the final population sizes of both macrophyte species with a marginal effect on growth rate of *L. minor* and the chlorophyll A production of *S. polyrhiza*, but overall these species were very tolerant of microcystin. The strong tolerance supports the potential use of these plants as bioremediators of cyanobacterial blooms. The differential impact of microcystin exposure discovered in single lineage models among genotypes indicates a potential for cyanotoxins to act as selective forces and reduce local macrophyte genetic diversity.

**Highlights:**

- Ecotoxicology often uses standard genotypes of plants in testing.
- We tested the influence of clonal variation in duckweeds on their response to common cyanotoxin, microcystin-LR.
- Microcystin impacts were often masked by genotypic variation in response.
- Results imply that genotype identity may be important to bioremediation and local evolutionary dynamics.

## 1. Introduction

Environmental chemical stress has long been recognized as a source of ecological change and, more recently, evolutionary change (Bickham 2011, Whitehead 2017, Brady et al. 2017). Such evolutionary changes are particularly pronounced in plant populations (Antonovics, 2006; Pitelka, 1988; Wu and Bradshaw, 1972). The demonstrated tolerance evolution has augmented interest in the use of plant bioremediation as anthropogenic influence is increasing the prevalence of both synthetic chemical stressors and naturally occurring toxins (Noyes et al., 2009; Schiedek et al., 2007). The increase of naturally occurring toxins such as allelochemicals, chemicals exuded by a plant or bacterium to decrease the success of competitors (Molisch 2001), can lead to complicated competition dynamics between toxin-producing organisms and toxin-receiving organisms (Driscoll et al., 2015; Lankau, 2009). Despite the ubiquitous presence of allelochemicals in both terrestrial and aquatic environments (Inderjit et al., 2008; Kalisz et al., 2021), the significance of increased allelopathy in aquatic environments and related bioremediation efforts has been relatively unexplored.

Cyanobacterial harmful algal blooms (CyanoHABs) are dense aggregates of cyanobacteria, a group of phytoplankton which produce allelochemicals known as cyanotoxins, that form in freshwater systems. Cyanotoxins are responsible for large economic losses, are detrimental to human health, and cause widespread ecological damage through the release of toxin as well as the creation of anoxic conditions within lakes (Cheung et al., 2013; Heisler et al., 2008). Furthermore, cyanotoxin release is anticipated to increase in both frequency and magnitude in response to eutrophication and climate change (Cheung et al., 2013).

While proposed mechanisms for controlling CyanoHABs have focused on abiotic factors (reviewed in: Fu et al., 2012; Nwankwegu et al., 2019), bioremediation may provide a safe and tractable method of harm reduction. Aquatic plants, hereafter macrophytes, have the potential to act as biocontrol agents for CyanoHABs. Macrophytes interact with cyanobacteria through competition for key nutrients and light by shading the water column (Leflaive and Ten-Hage, 2007). Sufficient competition may therefore reduce cyanobacteria to a lower population size incapable of producing harmful concentrations of cyanotoxins. Additionally, some macrophytes may relieve environmental stress on other community members through direct uptake of cyanotoxins (e. g., Leflaive and Ten-Hage, 2007; Mitrovic et al., 2005; Nimptsch et al., 2008).

Duckweeds (family: Lemnacaea), a family of small, floating macrophytes, have been proposed as bioremediators for a variety of chemical stressors because of their rapid clonal reproduction and ability to uptake numerous pollutants (Ekperusi et al., 2019; Landolt, 1986; Ziegler et al., 2016, 2015). The rapid clonal reproduction allows for rapid growth of macrophytes in short periods of time, making duckweeds ideal candidates to quickly colonize polluted regions. Given the ability of duckweeds to form dense mats along the water’s surface in short time periods, they can not only reduce light availability but also reduce the shared nutrient pool in the water column (Ceschin et al., 2020; van Gerven et al., 2015). In fact, duckweeds are currently used as bioremediators in both agricultural ponds and wastewater treatment centers due to their extraordinary abilities to rapidly reduce nutrient loads and ease of harvest (Ziegler et al., 2016). Furthermore, duckweeds are known to co-occur with CyanoHABs and uptake cyanotoxin in controlled experiments (Kaminski et al., 2014; Li et al., 2020; Nimptsch et al., 2008). Therefore, duckweeds may pose a stronger competitive pressure than rooted, submerged macrophytes while limiting toxin persistence.

However, standardized practices in ecotoxicological studies using duckweeds are often limited to a single genotype (ASTM International, 2022) or performed without regard to genotype identity (i.e., effects are averaged over a population). Although standardized practices benefit comparisons among stressor impacts within single species, duckweeds are known to possess intraspecific variation in a variety of traits including pollutant uptake, growth across nutrients levels, and photosynthetic ability under the stress regimes (e.g., Anneberg et al., 2023; Chen et al., 2020; Roubeau Dumont et al., 2019). Given the evidence of intraspecific variation to other chemical stressors, neglecting such variation may impact the efficacy of predicting duckweed-cyanobacteria interactions in CyanoHAB bioremediation. For example, cyanotoxin-induced damage to relevant photosynthetic processes may vary among genotypes of the same species. The toxin may act upon this pre-existing variability in sensitivity as a selection pressure and thus reduce macrophyte populations to only resilient plants, limiting overall genetic diversity and possibly altering stability to other stressors. Such natural selection for resilience could translate into direct management implications. Bioremediation efforts may fail by reducing the fitness of sensitive genotypes beyond thresholds of persistence or through the emergence of “cheater” genotypes if detoxification is a public good (Shibasaki and Mitri, 2020). Understanding intraspecific variation in cyanotoxin response is necessary to enact successful bioremediation plans.

To determine if interspecific or intraspecific variation in duckweed species response to a cosmopolitan cyanotoxin exist, we challenged fifteen genotypic lineages each for two duckweed species with ecologically relevant concentrations of microcystin-LR. We assessed biomarkers related to performance (total yield as biomass and photosynthetic potential as chlorophyll A concentrations) and fitness (total population size, growth rate, and reproduction via dormant individuals). Using these biomarkers, we establish, 1) the sensitivity of each species to cyanotoxin presence and 2) the intraspecific variation in response within each species.

## 2. Methods

### 2.1 Study organisms

We selected fifteen genotypes of two globally distributed duckweed species, *Lemna minor* and *Spirodela polyrhiza* (Landolt 1986, Armitage and Jones, 2020). Gentoypes were selected from collections originating in Pennsylvania, Connecticut, Michigan, and Wisconsin USA to encompass a large geographic range in the Laurentian Great Lakes Basin and Eastern United States across which CyanoHABs are known to occur. Genotypes in this study were previously confirmed by microsatellite markers (Kerstetter et al., 2023) and only one genotype from each collection site was used. We sterilized duckweeds of both species by submerging a frond (i.e., a single duckweed individual) in a 5% hypochlorite solution for approximately 10 seconds; fronds then floated in the hypochlorite solution for an additional five minutes. We then established axenic cultures from a single frond in sterile test tubes containing 10% Lemnaceae-nutrient media (described in Appenroth 2015), except for adjusting the stoichiometric molar ratio of total nitrogen to phosphorus to 14:1. By establishing cultures from a single frond, genotypic lineages were maintained. Before the experiment, we transferred fronds to common garden conditions to grow for three weeks and remove maternal effects. Monocultures of fifteen genotypes per species grew in individual 150 mL Erlenmeyer flasks with 25% Lemnaceae-nutrient media. We housed macrophytes in Conviron BDW40 Plant Growth Chamber (Conviron, ND, USA) maintained at 23.5°C, 50% relative humidity, 150 µMol m s^-1^ LED light output with a 16:8 h light:dark cycle for the duration of the common garden and subsequent sensitivity assays.

### 2.2 Sensitivity assays

After common garden cultivation, we transferred fronds to the sensitivity assay. The sensitivity assay challenged duckweed fronds with 25% Lemnaceae-nutrient media containing microcystin-LR (MilliPoreSigma, MA, USA, cat. no.: 101043-37-2). Microcystin-LR is the most common isomer in both occurrence and concentration of microcystin in cyanoHABs and therefore an appropriate representative (Corbel et al., 2014; Ettoumi et al., 2011). Using an isolate of microcystin rather than direct exposure to the toxin-producing cyanobacteria avoids confounding the effects of competition with the effects of the toxin itself. Microcystin-LR concentrations treatments included 0 µg/L (control), 5 µg/L, 10 µg/L, and 15 µg/L. Selected concentrations are realistic to CyanoHAB conditions to maintain ecological relevance of allelochemical stress (e.g., Mishra et al., 2021) while assessing a gradient of dose-responses. We replicated microcystin treatments ten times for each genetic lineage in randomized positions within six well plates containing 12.5 mL of media (N = 960). Each replicate was seeded with four sterilely transferred fronds on day 0. Every other day thereafter until the completion of exposures on day 12, we photographed each plate from above to monitor population sizes. We ended exposures at twelve days to avoid confounding effects of intraspecific competition at high densities.

Upon completion of exposures, we collected plant material from the wells to obtain final biomass and chlorophyll A (chlA) concentration. We freeze-dried biomass at -100°C for 48 hours. Afterwards, the dry mass (mg) was measured using a Mettler Toledo analytical balance (precision: 0.000 mg; Mettler Toledo, OH, USA). We extracted chlA as a proxy for photosynthetic potential from freeze-dried material following standard protocols for plants (Wetzel and Likens, 2000; Zhao et al., 2017). In brief, we placed freeze-dried and massed plant material in 2 mL Eppendorf tubes, pestled the tissue, and added 1500 µL of ice-cold 70% acetone. We covered the tubes in foil to avoid photobleaching and stored at 4°C for a minimum of 48 hours until the material was completely devoid of pigment. After extractions, absorbance was read from three technical replicates per sample at 730 nm, 665 nm, and 645 nm using an Epoch microplate spectrophotometer and Biotek Gen5 software (BioTek U.S., VT, USA). We averaged the three technical replicates for a representative measurement of the sample. Concentration was calculated using the following equation as in Wetzel and Likens (2000): chlA mg/L = 11.75(A665-A730)-1.31(A645-A730). Chlorophyll A concentrations standardized to the mass of duckweed were then calculated as follows: chlA mg/mg duckweed = chlA mg/L (0.0015 L)/mg of duckweed.

Images were processed using ImageJ software (U. S. National Institutes of Health, MD, USA) using the multipoint tool to select and count fronds in each well. We counted all visible, vegetative fronds. *Spirodela polyrhiza* produces dormant storage buds known as turions under certain conditions that may overwinter (Landolt, 1986). Turions were collected and counted at the completion of the assays rather than as a measure of population growth during image processing as turions often drift to the bottom of experimental units and would not be visible in photos.

### 2.3 Statistical analysis

We assessed differences in population biomass, chlA concentrations, final population size, and turion count by model comparison. We fit linear mixed effects models using the ‘lmer’ function to compare biomass and chlA and the ‘glmer’ function using a Poisson distribution for count data to compare final population size and turion count from the lme4 package in R (Bates et al., 2023). Due to differences in species body size, biomarkers often exhibited a bimodal distribution that was attributable to the species identification. Therefore, we fit models for each species separately. We used genetic lineage (random effect) and microcystin treatment (fixed effect) to predict biomarkers. For all models, concentration of microcystin (treatment) was denoted as a categorical rather than continuous variable. Treating discrete doses in a limited range of concentrations as a continuous variable is often inappropriate due to the likelihood of hormesis (non-linearity wherein low or intermediate stressors may be stimulating) in biomarker responses (reviewed in: Calabrese and Blain, 2009; e.g., Bibo et al., 2008; Machado et al., 2017). Likelihood ratio tests were completed between subsets of competing models to determine the improvement of model fit given by each variable in the global model. Due to statistical power limitations and to avoid singularity, models were also fit for singular genetic lineages to determine the possibility of unique, lineage-specific impacts of treatment rather than the inclusion of random slopes.

To assess population growth, we fit exponential growth models to each replicate and compared the resulting growth rate. Using the ‘nlme’ function from the nlme package (Pinheiro et al., 2023), we set starting parameter values for the initial population size to 4 and starting per capita growth rate to 0.2. Fit growth rates were then compared in linear mixed effects models as described above for other biomarkers. All statistics were completed using RStudio version 1.2.5042 (R Core Team, Version 4.2.0).

## 3. Results

### 3.1 Microcystin concentration did not have a strong effect on the biomass of most genetic lineages of either species

We found duckweed genotype strongly influenced the total biomass for both species (*p <* 0.0001). Despite microcystin treatment showing no impact on total biomass of *L. minor* or *S. polyrhiza* (*p* = 0.52 and *p* = 0.12 respectively), variation among the genotypes within each species is clear. Although many genetic lineages were unaffected by microcystin concentration, others showed distinctive patterns (Fig. 1). For example, a pronounced increase in biomass as toxin concentration increased occurred in *L. minor* genotype L (*p* = 0.052, single genotype ANOVA) and *S. polyrhiza* lineage C (*p* = 0.064, single genotype ANOVA) in contrast to other lineages such as *S. polyrhiza* lineage O that showed an increase only in the 10 µg/L concentration (Fig. 1).

**Figure 1:**
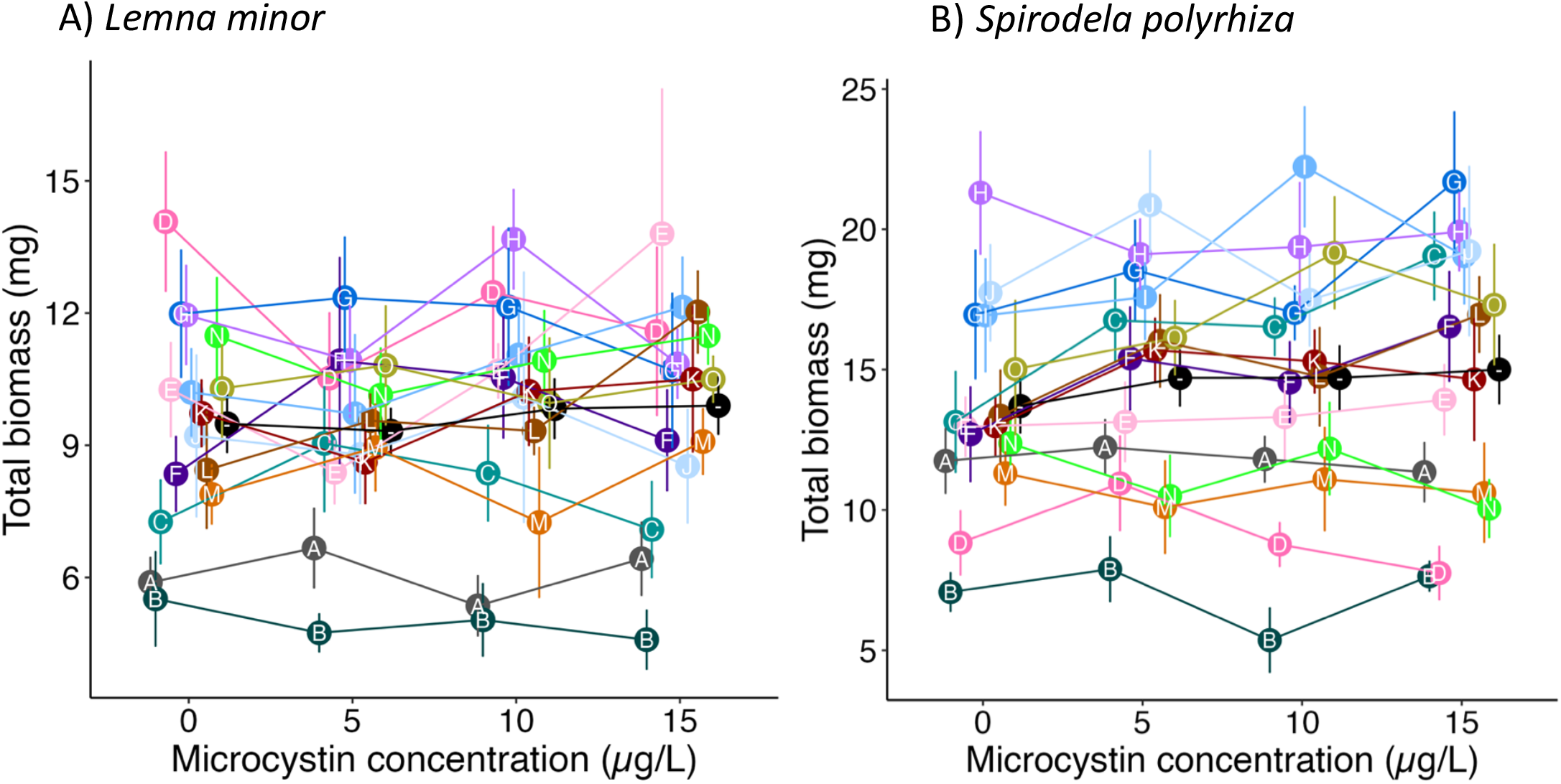
Total biomass (mg) at the completion of the experiment. Microcystin had a general stimulatory effect, though weak and not significant (estimated marginal means shown in black, labelled with “-”). Genotypic variation (genotypes labeled by letter, color) from this trend is apparent in both species. Each point represents the mean with standard error bars (n=10).

### 3.2 Photosynthetic activity of S. polyrhiza was more sensitive to microcystin concentration than L. minor

Chlorophyll A content per mg tissue was strongly influenced by genotype for both species (*L. minor, p* < 0.0001; *S. polyrhiza, p* = 0.012), but microcystin treatment impact was stronger on *S. polyrhiza* (*p* = 0.085, Fig. 2B) than on *L. minor* chlA content (*p* = 0.87; Figure 2A). Patterns among genotypes are not clear for *L. minor*, but *S. polyrhiza* genotypes generally decreased at the lowest and highest concentrations of microcystin, indicating nonlinearity of response to the dose concentration. This effect was pronounced in *S. polyrhiza* genotype A (*p* = 0.04, single genotype ANOVA) and genotype F (*p* = 0.08, single genotype ANOVA). Some *S. polyrhiza* genotypes deviated from this general pattern and instead showed a decrease across concentrations as in genotype C (*p* = 0.02, single genotype ANOVA) or relative insensitivity (Fig. 2B).

**Figure 2:**
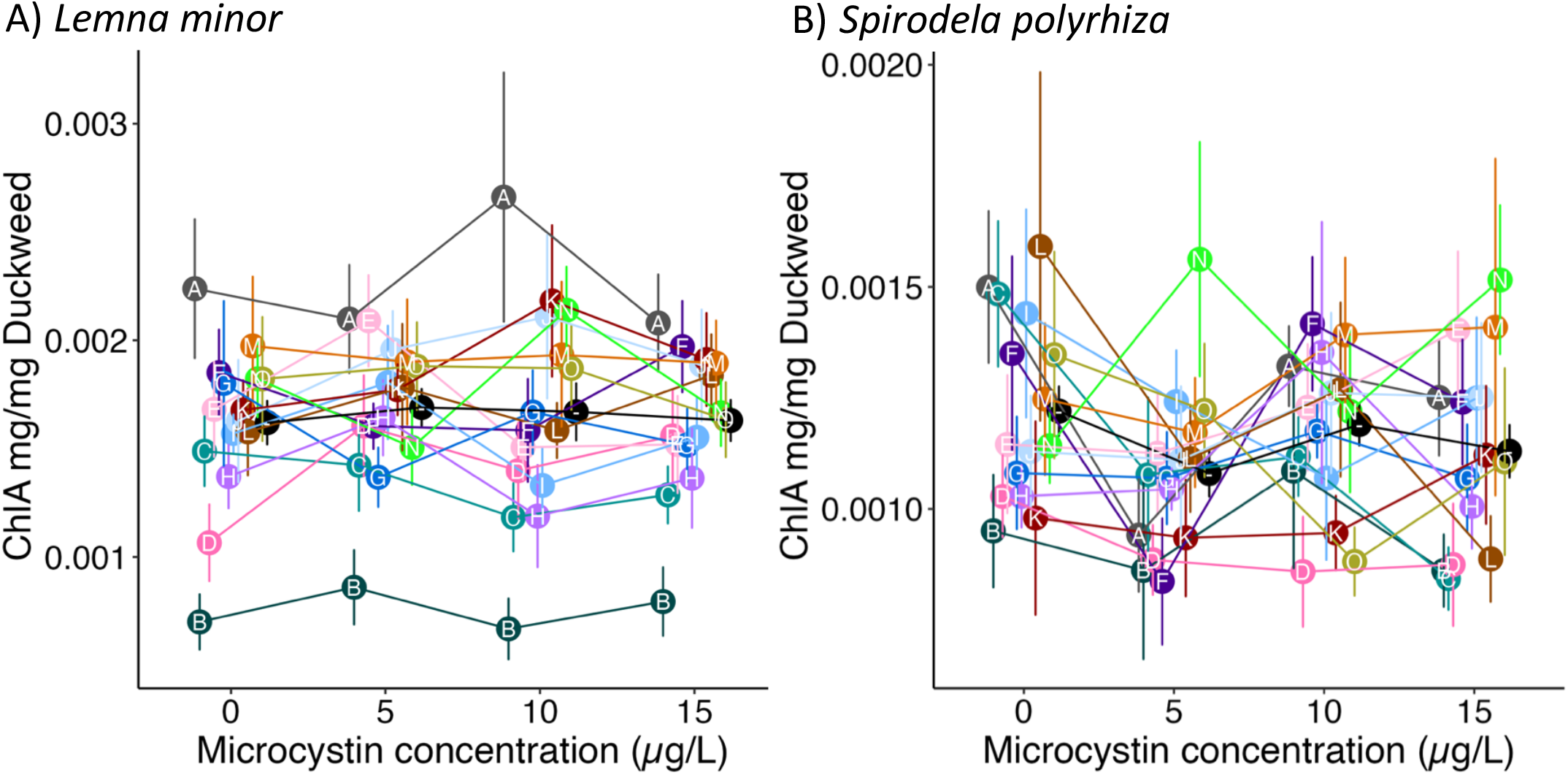
Chlorophyll A (chlA) production standardized to biomass. Microcystin had little impact on *Lemna minor* photosynthetic potential, but did show a decrease in *Spirodela polyrhiza* photosynthetic potential at the lowest concentration (estimated marginal means shown in black, labelled with -). Genotypic variation (genotypes labeled by letter, color) in *L. minor* may account for the lack of directional impact of microcystin concentration alone. Each point represents the mean with standard error bars (n=10).

### 3.3 Fitness metrics varied by genetic lineage and microcystin concentration

We assessed both the final frond number and exponential growth rate as well as turion production to determine fitness effects. The final number of fronds after twelve days of growth for both species was significantly influenced by genotype (*L. minor, p* < 0.0001; *S. polyrhiza, p* < 0.0001) and microcystin concentration for *L. minor* only (*L. minor, p* = 0.003; *S. polyrhiza, p* = 0.103; Fig. 3A and B respectively). Trends towards a higher population size with increased concentration appeared to be driven by two *L. minor* lineages, I and L (*p* = 0.001 and *p* < 0.001 respectively, single genotype ANOVA), and overall effect sizes of microcystin concentration only accounted for a difference of four fronds between the control and highest concentrations. In contrast, *L. minor* lineages B and C experienced a slight decrease in population size across the gradient (*p* = 0.03 and *p* = 0.03 respectively, single genotype ANOVA). Such differences in directionality emphasize the potential interaction between treatment and genotype. Effect sizes of microcystin treatment on *S. polyrhiza* were neglible, representing a single frond difference. Genotypes G and K initially increased at low concentrations and returned to control population sizes at higher microcystin concentrations (*p* < 0.001 for each, single genotype ANOVA), but other *S. polyrhiza* genetic lineages were insensitive (Fig. 3B).

**Figure 3:**
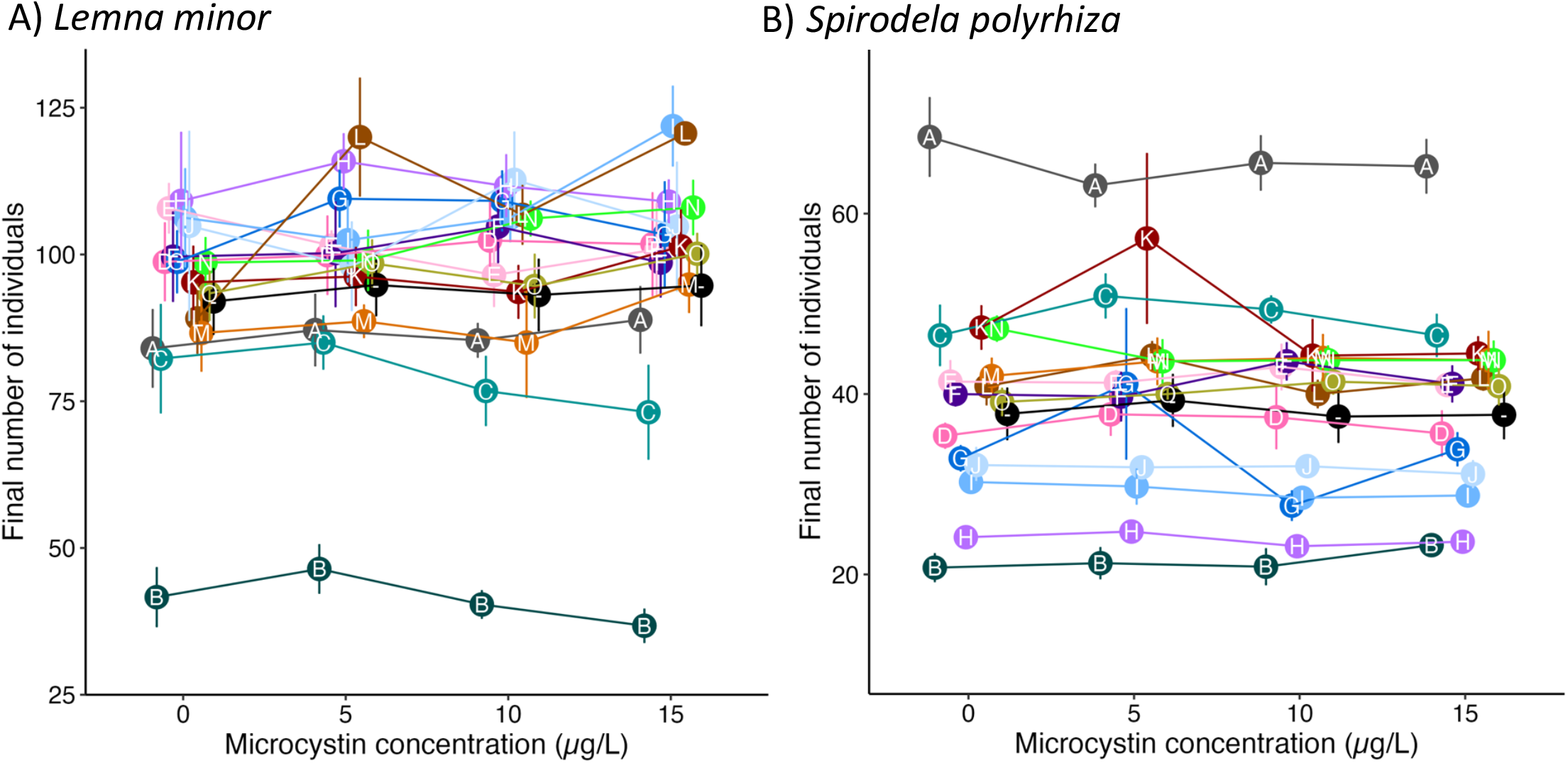
Final population size (number of individual duckweed fronds) at the end of experiments. Exposures were terminated after twelve days of population growth. Microcystin concentration showed a weak stimulatory effect in *Lemna minor* but not *Spirodela polyrhiza* (estimated marginal means shown in black, labelled with “-”). Despite this general trend, some genotypes (labeled by letter, color) were insensitive while others declined across the gradient. Each point represents the mean with standard error bars (n=10).

Despite the influence of microcystin exposure on population size, exponential growth rates were not affected. We fit exponential growth models to each well. Microcystin concentration was not found to be a significant predictor of either *L. minor* (*p* = 0.129) or *S. polyrhiza* (*p* = 0.479) exponential growth rates though patterns of weak increase are visible for *L. minor* (Fig. 4A). However, *S. polyrhiza* growth rates modestly increased at the 5 µg/L microcystin (Fig. 4B). For example, *S. polyrhiza* genotypes K and G all decreased in growth at the 10 µg/L concentration while the pink genotype increased at this level (Fig. 4B).

**Figure 4:**
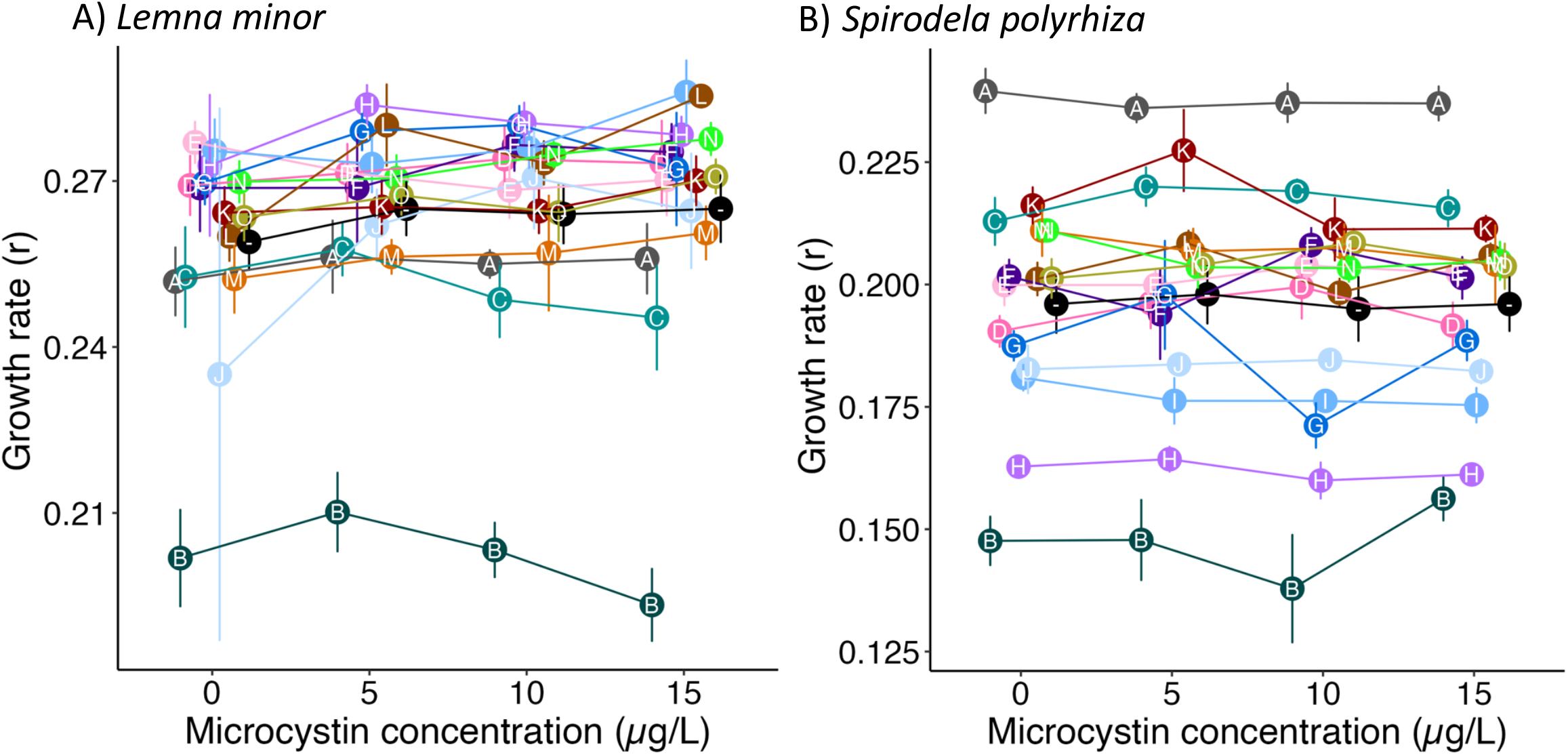
Exponential growth rate across a microcystin gradient. We fit exponential growth models to frond abundance data across the twelve-day exposure period. Microcystin concentrations led to a modest increase in growth rate (estimated marginal means in black, labelled with -) that is more apparent for *Lemna minor* than *Spirodela polyrhiza*. Genotypes (labeled by letter, color) varied with some sensitive lines declining. Each point represents the mean with standard error bars (n=10).

Turion production in *S. polyrhiza* was attributable to both genetic lineage (*p <* 0.0001) and microcystin concentration (*p* = 0.0002). As concentration increased, turion production mildly increased (Fig. 5). This trend was most pronounced in lineage G (single lineage GLM, *p* = 0.0002). Some deviance in trend was noted among the genetic lineages. For example, lineage D showed a U-shaped response with an initial increase at low concentrations of microcystin and a decrease in the highest concentration relative to the control (single lineage GLM, *p* = 0.0001) similar to lineage N for which intermediate levels showed a stimulatory effect (single lineage GLM, *p* = 0.001).

**Figure 5:**
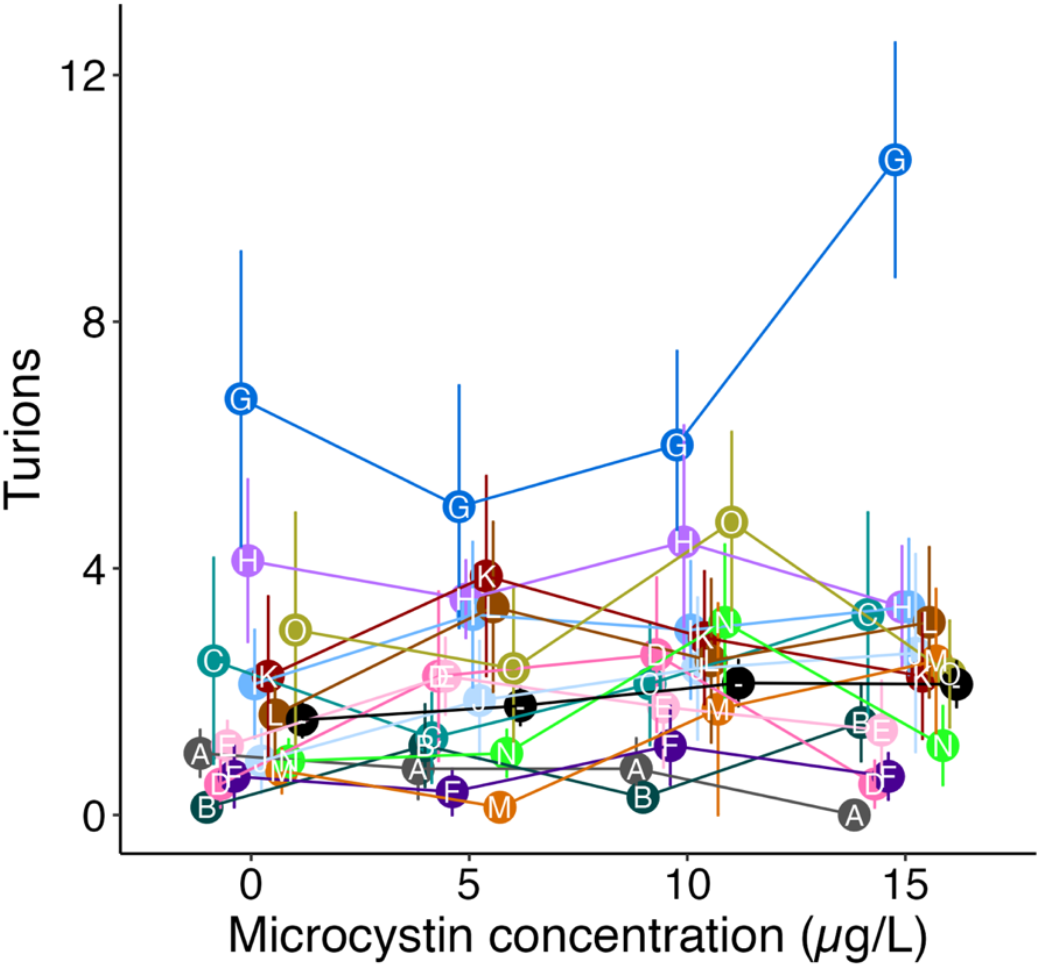
Turion production across a microcystin gradient. The production of dormant buds, turions, increased with microcystin exposure (*p* = 0.0002; estimated marginal means shown in black). The magnitude of effect varied by genotype (shown in colors, labelled by letter.

## 4. Discussion

We established both intraspecific and interspecific variation in macrophyte response to cyanotoxin stress. While microcystin responses resulted in relatively small impacts on duckweed biomarkers in comparison to the impact of genotype alone, genotypes often displayed several patterns of response within biomarker. Population level assessments may inappropriately conclude that cyanotoxin has no effect on selected species despite unique trends within genetic lineages. The sensitivity of macrophyte traits to microcystin exposure demonstrated differential productivity (chlA concentration and biomass) and fitness (final population size, growth rate, and turions) within species with stimulatory effects more pronounced in *L. minor* lineages than their *S. polyrhiza* counterparts for which nonlinear responses were more common. These differences in sensitivity highlight the importance of not only optimal species selection in bioremediation but also of optimal genotype selection. Furthermore, the intraspecific variation demonstrates that microcystin and similar cyanotoxins may act as a selective force on macrophyte tolerance.

Duckweeds have been used as bioremediation agents for a variety of chemical stressors (Bassuney and Tawfik, 2017; Ekperusi et al., 2019; Liu et al., 2020; Ziegler et al., 2016). However, standardized practices for testing bioremediation potential and ecotoxicological effects of stresses reduce intraspecific variation to single genetic lineages (ASTM International, 2022). Our results demonstrate that this reductionist approach may limit our ability to predict how duckweeds will perform as bioremediation agents in natural or constructed populations. If a single genotype is found to be highly sensitive to a cyanotoxin, we may incorrectly assume that the entire species is inappropriate to use as a bioremediator. Furthermore, cyanotoxins did not have linear impacts within both species. Such phenomena of low dose stimulation or general U-shaped, non-linear responses to toxin and toxicant exposure is common among plants and across aquatic taxa (Agathokleous et al., 2021; Calabrese and Blain, 2009). This suggests that the genetic makeup of duckweed populations differentially affect population-level responses to harmful algal blooms. Such interactions stress the importance of screening multiple genotypes for bioremediation.

When considered at the population level, however, both species were relatively tolerant of cyanotoxin exposure at ecologically relevant levels. In tandem with previous evidence of high uptake rates of cyanotoxin by *L. minor* and other duckweed species (Kaminski et al., 2014; Li et al., 2020; Nimptsch et al., 2008), our results confirm that duckweed species may be used as successful bioremediators. Both species contained genotypes that were capable of suriviving and maintaining productivity similar to that of control conditions. We do note that the fifteen genotypes tested for each species were collected throughout the Northern United States where environmental history of blooms is likely. It is possible that the general tolerance seen in our experiment is related to long-term adaptation. The use of duckweeds as bioremediators of cyanobacterial blooms may be enhanced through selecting genotypes stimulated in growth by toxin presence. Such genotypes may have higher population level uptake rates or increase competitive pressure on cyanobacteria through additional shading and nutrient removal to limit bloom size.

Beyond management implications, we demonstrated microcystin may act as a selective agent through the differential performance of genotypes when considered at the within genotype level. We found that while many genotypes of both species slightly increased exponential growth rate during microcystin exposure, others’ growth rates were not stimulated or even depressed. While our study addresses the exponential growth rate in monoculture, these differences may lead to changes in genotypic frequency over time. Previous studies of allelochemicals as selective agents have largely addressed terrestrial environments (Inderjit et al., 2008; Lankau, 2009; Lankau and Strauss, 2007) with little attention to aquatic environments. In cases where allelochemicals are known selective agents, eco-evolutionary dynamics may ensue. For example, the release of allelochemical may select for tolerance or resilience in competing species (i.e., select for genotypes that maintain fitness under stress). An arms race may be established between the toxin producer and recipient species.

Thus, the dual ecological and evolutionary forces caused by microcystin demonstrate the CyanoHABs may be used as a model system to test eco-evolutionary hypotheses. The system is amenable for highly replicated laboratory studies of short duration and may be valuable to testing a variety of hypotheses on the evolution of toxigenicity (Driscoll et al., 2015). Microcystin, the cyanotoxin assessed in this study, is created through a heritable genetic mechanism that is found in only some individuals within cyanobacteria populations (Rantala et al., 2004; Wilson et al., 2005). Therefore, both evolution of cyanobacterial toxigenicity and ecological changes in a CyanoHAB can be tested while exploring eco-evolutionary dynamics of community members interacting with cyanotoxins. Using the CyanoHAB system, we can address topics such as when toxigenicity and/or cyclical eco-evolutionary dynamics are each favored, and how toxigenicity can alter system stability. Augmenting our understanding of these dynamics can help us improve predictive management tools to limit human toxin exposure and bloom formation.

## 5. Conclusions

Our study indicates the need for consideration of genotypes and eco-evolutionary dynamics in bioremediation of CyanoHABs. Microcystin effects were often masked by conflicting directional responses among genotypes. Therefore, greater attention must be placed on not only population level assessments of macrophyte remediators but also the genotypic background of such populations. Furthermore, differential fitness of macrophyte genotypes allows the CyanoHAB system to provide insights on broader questions on the evolution of toxigenicity and ensuing eco-evolutionary dynamics.

## Acknowledgments

We would like to acknowledge the assistance of Jae Kerstetter and Audrey Burr for duckweed colony maintenance and assistance in daily monitoring. We would also like to thank undergraduate researchers of the Turcotte lab for their assistance in data processing. This work was partially supported by an NSF grant (#1935410) to M.M.T.

## Notes

### Competing Interest Statement

The authors have declared no competing interest.

## Literature Cited

Agathokleous, E., Barceló, D., Fatta-Kassinos, D., Moore, M.N., Calabrese, E.J., 2021. Contaminants of emerging concern and aquatic organisms: the need to consider hormetic responses in effect evaluations. Water Emerg. Contam. Nanoplastics 1, 2. https://doi.org/10.20517/WECN.2021.01

Anneberg, T.J., O’Neill, E.M., Ashman, T.L., Turcotte, M.M., 2023. Polyploidy impacts population growth and competition with diploids: multigenerational experiments reveal key life-history trade-offs. New Phytol. 238, 1294–1304. https://doi.org/10.1111/NPH.18794

Antonovics, J., 2006. Evolution in closely adjacent plant populations X: long-term persistence of prereproductive isolation at a mine boundary. Hered. 2006 971 97, 33–37. https://doi.org/10.1038/sj.hdy.6800835

Appenroth, K.J., 2015. Media for in vitro cultivation of duckweed. Duckweed Forum 3, 180–186. Available online at: http://www.ruduckweed.org/uploads/1/0/8/9/10896289/iscdra_issue11-2015-11_final.pdf.

Armitage, D.W., Jones, S.E., 2020. Coexistence barriers confine the poleward range of a globally distributed plant. Ecol. Lett. 23, 1838–1848. https://doi.org/10.1111/ELE.13612

ASTM International. 2022. Designation E1415-22.Standard guide for conducting static toxicity tests with Lemna Gibba G3. https://www.astm.org

Bassuney, D., Tawfik, A., 2017. Baffled duckweed pond system for treatment of agricultural drainage water containing pharmaceuticals. Int. J. Phytoremediation 19, 774–780. https://doi.org/10.1080/15226514.2017.1284756

Bates, D., Maechler, M., Bolker, B., Walker, S., Christensen, R.H.B., Singmann, H., Dai, B., Scheipl, F., Grothendieck, G., Green, P., Fox, J., Bauer, A., Krivitsky, P.N., 2023. Package “lme4.”

Bibo, L., Yan, G., Bangding, X., Jiantong, L., Yongding, L., 2008. A laboratory study on risk assessment of microcystin-RR in cropland. J. Environ. Manage. 86, 566–574. https://doi.org/10.1016/J.JENVMAN.2006.12.040

Calabrese, E.J., Blain, R.B., 2009. Hormesis and plant biology. Environ. Pollut. 157, 42–48. https://doi.org/10.1016/J.ENVPOL.2008.07.028

Ceschin, S., Crescenzi, M., Iannelli, M.A., 2020. Phytoremediation potential of the duckweeds Lemna minuta and Lemna minor to remove nutrients from treated waters. Environ. Sci. Pollut. Res. 27, 15806–15814. https://doi.org/10.1007/S11356-020-08045-3/FIGURES/6

Chen, Daoqian, Zhang, H., Wang, Q., Shao, M., Li, X., Chen, Dongmei, Zeng, R., Song, Y., 2020. Intraspecific variations in cadmium tolerance and phytoaccumulation in giant duckweed (Spirodela polyrhiza). J. Hazard. Mater. 395, 122672. https://doi.org/10.1016/J.JHAZMAT.2020.122672

Cheung, M.Y., Liang, S., Lee, J., 2013. Toxin-producing cyanobacteria in freshwater: A Review of the problems, impact on drinking water safety, and efforts for protecting public health. J. Microbiol. 51, 1–10. https://doi.org/10.1007/s12275-013-2549-3

Corbel, S., Mougin, C., Bouaïcha, N., 2014. Cyanobacterial toxins: Modes of actions, fate in aquatic and soil ecosystems, phytotoxicity and bioaccumulation in agricultural crops. Chemosphere. https://doi.org/10.1016/j.chemosphere.2013.07.056

Driscoll, W.W., Hackett, J.D., Egis Ferri Ere, R., 2015. Eco-evolutionary feedbacks between private and public goods: evidence from toxic algal blooms. https://doi.org/10.1111/ele.12533

Ekperusi, A.O., Sikoki, F.D., Nwachukwu, E.O., 2019. Application of common duckweed (Lemna minor) in phytoremediation of chemicals in the environment: State and future perspective. Chemosphere 223, 285–309. https://doi.org/10.1016/J.CHEMOSPHERE.2019.02.025

Ettoumi, A., Khalloufi, F. El, Ghazali, I. El, Oudra, B., Amrani, A., Nasri, H., Bouaïcha, N., 2011. Bioaccumulation of cyanobacterial toxins in aquatic organisms and its consequences for public health, in: Kattel, G. (Ed.), Zooplankton and Phytoplankton: Types, Characteristics and Ecology. Nova Science Publishers Inc., New York, pp. 1–34. https://doi.org/10.13140/2.1.1959.5044

Fu, F.X., Tatters, A.O., Hutchins, D.A., 2012. Global change and the future of harmful algal blooms in the ocean. Mar. Ecol. Prog. Ser. 470, 207–233. https://doi.org/10.3354/MEPS10047

Heisler, J., Glibert, P.M., Burkholder, J.M., Anderson, D.M., Cochlan, W., Dennison, W.C., Dortch, Q., Gobler, C.J., Heil, C.A., Humphries, E., Lewitus, A., Magnien, R., Marshall, H.G., Sellner, K., Stockwell, D.A., Stoecker, D.K., Suddleson, M., 2008. Eutrophication and harmful algal blooms: A scientific consensus. Harmful Algae 8, 3–13. https://doi.org/10.1016/j.hal.2008.08.006

Inderjit Seastedt, T.R., Callaway, R.M., Pollock, J.L., Kaur, J., 2008. Allelopathy and plant invasions: Traditional, congeneric, and bio-geographical approaches. Biol. Invasions 10, 875–890. https://doi.org/10.1007/s10530-008-9239-9

Kalisz, S., Kivlin, S.N., Bialic-Murphy, L., 2021. Allelopathy is pervasive in invasive plants. Biol. Invasions 23, 367–371. https://doi.org/10.1007/s10530-020-02383-6

Kaminski, A., Bober, B., Chrapusta, E., Bialczyk, J., 2014. Phytoremediation of anatoxin-a by aquatic macrophyte Lemna trisulca. Chemosphere 112, 305–310. https://doi.org/10.1016/j.chemosphere.2014.04.064

Kerstetter, J.E., Reid, A.L., Armstrong, J.T., Zallek, T.A., Hobble, T.T., Turcotte, M.M., 2023. Characterization of microsatellite markers for the duckweed Spirodela polyrhiza and Lemna minor tested on samples from Europe and the United States of America.: Spirodela polyrhiza and Lemna minor microsatellites. Genet. Resour. 4, 46–55. https://doi.org/10.46265/GENRESJ.ALFV3636

Landolt, E., 1986. Biosystematic investigations in the family of duckweeds (Lemnaceae). Vol. 2.The family of Lemnaceae-A monographic study. Part 1 of the monograph: Morphology; karyology; ecology; geographic distribution; systematic position; nomenclature; descriptions. Publications of the Geobotanical Institute, ETH, Zurich Switzerland.

Lankau, R.A., 2009. Genetic variation promotes long-term coexistence of Brassica nigra and its competitors 174. https://doi.org/10.1086/600083

Lankau, R.A., Strauss, S.Y., 2007. Mutual feedbacks maintain both genetic and species diversity in a plant community. Science 317, 1561–1563. https://doi.org/10.1126/SCIENCE.1147455

Leflaive, J., Ten-Hage, L., 2007. Algal and cyanobacterial secondary metabolites in freshwaters: A comparison of allelopathic compounds and toxins. Freshw. Biol. https://doi.org/10.1111/j.1365-2427.2006.01689.x

Li, S., Le, S., Li, G., Luo, M., Wang, R., Zhao, Y., 2020. Bioremediation of Landoltia punctata to Microcystis aeruginosa contaminated waters. Water 2020, Vol. 12, Page 1764 12, 1764. https://doi.org/10.3390/W12061764

Liu, Y., Xu, H., Yu, C., Zhou, G., 2020. Multifaceted roles of duckweed in aquatic phytoremediation and bioproducts synthesis. GCB Bioenergy. 13, 70–82. https://doi.org/10.1111/gcbb.12747

Machado, J., Campos, A., Vasconcelos, V., Freitas, M., 2017. Effects of microcystin-LR and cylindrospermopsin on plant-soil systems: A review of their relevance for agricultural plant quality and public health. Environ. Res. 153, 191–204. https://doi.org/10.1016/J.ENVRES.2016.09.015

Mishra, S., Stumpf, R.P., Schaeffer, B., Werdell, P.J., Loftin, K.A., Meredith, A., 2021. Evaluation of a satellite-based cyanobacteria bloom detection algorithm using field-measured microcystin data. Sci. Total Environ. 774, 145462. https://doi.org/10.1016/J.SCITOTENV.2021.145462

Mitrovic, S.M., Allis, O., Furey, A., James, K.J., 2005. Bioaccumulation and harmful effects of microcystin-LR in the aquatic plants Lemna minor and Wolffia arrhiza and the filamentous alga Chladophora fracta. Ecotoxicol. Environ. Saf. 61, 345–352. https://doi.org/10.1016/j.ecoenv.2004.11.003

Nimptsch, J., Wiegand, C., Pflugmacher, S., 2008. Cyanobacterial toxin elimination via bioaccumulation of MC-LR in aquatic macrophytes: An application of the “Green Liver Concept”; Environ. Sci. Technol. 42, 8552–8557. https://doi.org/10.1021/es8010404

Noyes, P.D., McElwee, M.K., Miller, H.D., Clark, B.W., Van Tiem, L.A., Walcott, K.C., Erwin, K.N., Levin, E.D., 2009. The toxicology of climate change: environmental contaminants in a warming world. Environ. Int. 35, 971–986. https://doi.org/10.1016/J.ENVINT.2009.02.006

Nwankwegu, A.S., Li, Y., Huang, Y., Wei, J., Norgbey, E., Sarpong, L., Lai, Q., Wang, K., 2019. Harmful algal blooms under changing climate and constantly increasing anthropogenic actions: the review of management implications. 3 Biotech 9. https://doi.org/10.1007/S13205-019-1976-1

Pinheiro, J., Bates, D., DebRoy, S., Sarkar, D., Heisterkamp, S., Van Willigen, B., Ranke, J., R Core Team, 2023. Package “nlme.”

Pitelka, L.F., 1988. Evolutionary responses of plants to anthropogenic pollutants. Trends Ecol. Evol. 3, 233–236. https://doi.org/10.1016/0169-5347(88)90165-6

Rantala, A., Fewer, D.P., Hisbergues, M., Rouhiainen, L., Vaitomaa, J., Börner, T., Sivonen, K., 2004. Phylogenetic evidence for the early evolution of microcystin synthesis. Proc. Natl. Acad. Sci. 101, 568–573. https://doi.org/10.1073/PNAS.0304489101

Roubeau Dumont, E., Larue, C., Lorber, S., Gryta, H., Billoir, E., Gross, E.M., Elger, A., 2019. Does intraspecific variability matter in ecological risk assessment? Investigation of genotypic variations in three macrophyte species exposed to copper. Aquat. Toxicol. 211, 29–37. https://doi.org/10.1016/J.AQUATOX.2019.03.012

Schiedek, D., Sundelin, B., Readman, J.W., Macdonald, R.W., 2007. Interactions between climate change and contaminants. Mar. Pollut. Bull. 54, 1845–1856. https://doi.org/10.1016/J.MARPOLBUL.2007.09.020

Shibasaki, S., Mitri, S., 2020. Controlling evolutionary dynamics to optimize microbial bioremediation. Evol. Appl. 13, 2460–2471. https://doi.org/10.1111/EVA.13050

van Gerven, L.P.A., de Klein, J.J.M., Gerla, D.J., Kooi, B.W., Kuiper, J.J., Mooij, W.M., 2015. Competition for light and nutrients in layered communities of aquatic plants. Am. Nat. 186, 72–83. https://doi.org/10.1086/681620

Wetzel, R.G., Likens, G.E., 2000. Limnological Analyses, 3rd ed, Limnological Analyses. Springer New York, New York. https://doi.org/10.1007/978-1-4757-3250-4

Wilson, Alan E, Sarnelle, Orlando, Neilan, B.A., Salmon, T.P., Gehringer, M.M., Hay, M.E., Raikow, D.F., Hamilton, S.K., Oceanogr, L., 2005. Genetic variation of the bloom-forming cyanobacterium Microcystis aeruginosa within and among lakes: Implications for harmful algal blooms. Appl. Environ. Microbiol. 71, 6126–6133. https://doi.org/10.1128/AEM.71.10.6126-6133.2005

Wu, L., Bradshaw, A.D., 1972. Aerial pollution and the rapid evolution of copper tolerance. Nat. 1972 2385360 238, 167–169. https://doi.org/10.1038/238167a0

Zhao, Z., Shi, H., Kang, X., Liu, C., Chen, L., Liang, X., Jin, L., 2017. Inter- and intra-specific competition of duckweed under multiple heavy metal contaminated water. Aquat. Toxicol. 192, 216–223. https://doi.org/10.1016/j.aquatox.2017.09.023

Ziegler, P., Adelmann, K., Zimmer, S., Schmidt, C., Appenroth, K.-J., 2015. Relative in vitro growth rates of duckweeds (Lemnaceae) - the most rapidly growing higher plants. Plant Biol. 17, 33–41. https://doi.org/10.1111/plb.12184

Ziegler, P., Sree, K.S., Appenroth, K.J., 2016. Duckweeds for water remediation and toxicity testing: Toxicol. Environ. Chem. 98, 1127–1154. https://doi.org/10.1080/02772248.2015.1094701

